# Linking Enlarged Choroid Plexus with Plasma Analyte and Structural Phenotypes in Clinical High Risk for Psychosis: A Multisite Neuroimaging Study

**DOI:** 10.1101/2022.10.28.514160

**Authors:** Deepthi Bannai, Martin Reuter, Rachal Hegde, Dung Hoang, Iniya Adhan, Swetha Gandu, Sovannarath Pong, Nick Raymond, Victor Zeng, Yoonho Chung, George He, Daqiang Sun, Theo G.M. van Erp, Jean Addington, Carrie E. Bearden, Kristin Cadenhead, Barbara Cornblatt, Daniel H. Mathalon, Thomas McGlashan, Clark Jeffries, William Stone, Ming Tsuang, Elaine Walker, Scott W. Woods, Tyrone D. Cannon, Diana Perkins, Matcheri Keshavan, Paulo Lizano

## Abstract

**Background:** Choroid plexus (ChP) enlargement exists in first-episode and chronic psychosis, but whether enlargement occurs before psychosis onset is unknown. This study investigated whether ChP volume is enlarged in individuals with clinical high-risk (CHR) for psychosis and whether these changes are related to clinical, neuroanatomical, and plasma analytes.

**Methods:** Clinical and neuroimaging data from the North American Prodrome Longitudinal Study 2 (NAPLS2) was used for analysis. 509 participants (169 controls, 340 CHR) were recruited. Conversion status was determined after 2-years of follow-up, with 36 psychosis converters. The lateral ventricle ChP was manually segmented from baseline scans. A subsample of 31 controls and 53 CHR had plasma analyte and neuroimaging data.

**Results:** Compared to controls, CHR (*d=0*.*23, p=0*.*017*) and non-converters *(d=0*.*22, p=0*.*03)* demonstrated higher ChP volumes, but not in converters. In CHR, greater ChP volume correlated with lower cortical *(r=-0*.*22, p<0*.*001)*, subcortical gray matter *(r=-0*.*21, p<0*.*001)*, and total white matter volume *(r=-0*.*28,p<0*.*001)*, as well as larger lateral ventricle volume *(r=0*.*63,p<0*.*001)*. Greater ChP volume correlated with makers functionally associated with the lateral ventricle ChP in CHR [CCL1 *(r=-0*.*30, p=0*.*035)*, ICAM1 *(r=0*.*33, p=0*.*02)]*, converters [IL1β *(r=0*.*66, p=0*.*004*)], and non-converters [BMP6 *(r=-0*.*96, p<0*.*001)*, CALB1 *(r=-0*.*98, p<0*.*001)*, ICAM1 *(r=0*.*80, p=0*.*003)*, SELE *(r=0*.*59, p=0*.*026)*, SHBG *(r=0*.*99, p<0*.*001)*, TNFRSF10C *(r=0*.*78, p=0*.*001)*].

**Conclusions:** CHR and non-converters demonstrated significantly larger ChP volumes compared to controls. Enlarged ChP was associated with neuroanatomical alterations and analyte markers functionally associated with the ChP. These findings suggest that the ChP may be a key explanatory biomarker in CHR for psychosis.

## Introduction

The choroid plexus (ChP) projects into the four cerebral ventricles and is the principal source of cerebrospinal fluid (CSF). Together, the ChP and CSF play an important role in brain physiology^1,2^, buoyancy^3^, and homeostasis by providing physical, enzymatic, and immunological barriers to the brain^4^. Neuroimaging studies have observed ChP changes with aging^5–7^ and across various neurodevelopmental^8–10^ and brain disorders^11–13^, suggesting that the ChP may play a key role in brain disorders^14^. Despite growing evidence implicating the ChP in neurodevelopment and brain disorders, it has not been the focus of commonly used neuroimaging tools, which causes it to be poorly segmented, mislabeled, and incorrectly quantified^15^. Therefore, there is a critical need to elucidate the role of the ChP in brain disorders using more accurate segmentation of the ChP.

The earliest observations of ChP morphological changes in schizophrenia and bipolar disorders occurred in the early 20^th^ century, where cystic formations, fibrosis, lipid deposits, and hypersecretory phenotypes were found in ChP epithelial cells, as well as endothelial cell degradation^16–18^. We previously reported on ChP enlargement in a large sample of patients with schizophrenia, schizoaffective, and psychotic bipolar disorder compared to healthy controls^19^. Larger ChP volume was associated with worse overall cognition, but no correlations with clinical measures were identified. A link between ChP volume and brain structure was established, with greater ChP volume relating to smaller gray matter and subcortical volume, larger ventricular volume, and lower white matter microstructure. Enlarged ChP volume correlated with higher peripheral levels of interleukin-6, suggesting that inflammation may play a role in ChP structural changes observed in psychosis and inflammation-meditated CSF hypersecretion^20,21^. These findings were replicated in a study in first-episode schizophrenia, where the greater ChP volume observed also correlated with higher allostatic load (indexed by subclinical cardiovascular, metabolic, neuroendocrine, and immune markers)^22^. Another study reported larger ChP volume in schizophrenia patients with orofacial tardive dyskinesia and identified associations with elevated N-methyl-D-aspartate receptor antibody levels^23^. Taken together, these results suggest that the ChP may be an important structure in mechanisms underlying psychosis pathology. However, no studies to date have investigated ChP volumes in a population at clinical high risk (CHR) for psychosis to determine whether ChP enlargement is associated with future psychosis risk.

The results from these neuroimaging studies are promising; however, the use of FreeSurfer for ChP segmentation limits their interpretation, primarily due to poor ChP segmentation by FreeSurfer^15^. Recently, four tools have been developed to provide better ChP parcellation: Gaussian Mixture Model (GMM; Alzheimer’s)^24^, optimized 3D U-Net^25^ (aging healthy individuals), 2 step 3D U-Net^26^ (Multiple Sclerosis and healthy controls), and an axial multi-layer perceptron (Axial-MLP 8; Multiple Sclerosis and healthy controls)^27^. While robust, each technique only provides adequate segmentation performance, reflected in average Dice Coefficients (DC) of 64 for GMM, 71 for Axial-MLP8, 73 for 3D U-Net, and 72 for 2 step 3D-U-Net. Therefore, in this study, we aim to analyze ChP volume in individuals at CHR for psychosis using manually segmented lateral ventricle ChP from the North American Prodrome Longitudinal Study 2 (NAPLS2) and compare these manually-segmented values to two techniques previously used to study ChP volume in neuropsychiatric disorders: FreeSurfer 6.0 and GMM. We further aim to understand whether greater ChP volume is related to clinical, neuroimaging, and plasma analyte measures. We hypothesized that the CHR group will demonstrate ChP enlargement compared to healthy controls (HC) and that converters to psychosis will have higher volumes. Finally, we hypothesized that greater ChP volumes in CHR would be associated with worse cognition, global brain volume reduction, and higher plasma analyte levels.

## Methods

### Participants

Participants were part of NAPLS2, a prospective multisite study analyzing psychosis predictors in individuals 12-35 years of age meeting Criteria of Psychosis Risk Syndromes^28^. Institutional Review Board approval was obtained from all sites^29^. Participants included 521 individuals (175 HC and 346 CHR) who had completed T1 MRI scans. Each participant provided written consent and parents/guardians provided consent for minors with minors providing assent. Exclusion criteria included current or lifetime Axis I psychotic disorder, IQ<70, past history of a central nervous system disorder, substance dependence within the past 6 months, and prodromal symptoms caused by an Axis I disorders. Non-psychotic DSM-IV disorders were not exclusionary (e.g. substance abuse disorder, anxiety disorders, or Axis II disorders) if they did not account for psychosis-risk symptoms. Exclusionary criteria for HC included meeting psychosis-risk syndrome criteria, any current or past psychotic disorder or Cluster A personality disorder, first-degree family history of any psychotic disorder or any disorder involving psychotic symptoms, and currently using psychotropic medications. Symptoms were assessed using the Scale of Psychosis-risk Symptoms (SOPS)^29,30^. The Measurement and Treatment Research to Improve Cognition in Schizophrenia (MATRICS)^31–33^ and Global Assessment of Functioning (GAF)^34^ measures were gathered to evaluate cognition and functioning, respectively. Social cognition was tested using The Awareness of Social Inference Test (TASIT)^35^. A questionnaire documenting trauma from before the age of 16 was collected. Conversion status was determined up to 24-month-follow-up or at the point of conversion to psychosis for either a psychotic illness or bipolar disorder (bipolar 1/2, cyclothymic, or not-otherwise-specified). Substance use status was categorized by alcohol, amphetamine, cannabis, cocaine, hallucinogen, or polysubstance use. Illness duration categories (short or long) were determined by median split (30.9 weeks). Antipsychotic status and average daily chlorpromazine (CPZ) equivalents were collected^36^. One HC was dropped due to conversion to psychosis. Lastly, 5 participants were dropped due to missing diagnostic information.

### MRI data

Subjects had baseline MRI scans that were free of visually detectable artifacts. A detailed explanation of scanner parameters and site harmonization was detailed previously^37^. Five sites used Siemens 3T with a 12-channel head coil (Emory, Harvard, Yale, University of California Los Angeles, and University of North Carolina) and 3 used GE 3T with an 8-channel head coil (Zucker Hillside, University of California San Diego, and University of Calgary). Images were acquired in the sagittal plane with a 1×1×1.2mm resolution. Siemens devices used an MPRAGE sequence with a 256×240×176mm field of view, TR/TE/T1=2300/2.91/900ms, and a 9-degree flip angle. GE scanners used an IR-SPGR sequence with a 26cm field of view, TR/TE/TI=7.0/minimum full/400ms, and an 8-degree flip angle.

### Choroid plexus segmentation

T_1_-MPRAGE images were processed through FreeSurfer v6.0^38^. Upon visual inspection, FreeSurfer lateral ventricle ChP maps were found to segment ∼50% of the structure (Figure 1A,D,G,J). Thus, a Gaussian Mixture Model (GMM), previously developed for ChP analysis in Alzheimer’s disease, was used to segment ChP^24^. However, GMM ChP parcellations were found to be inaccurate (Figure 1B,E,H,K) and we decided to manually-segment the ChP for all scans. 3D Slicer v4.11 was used for manual segmentation by 8 trained raters (IA, DB, SG, RH, SP, NR, AZ) in a first pass (our detailed protocol can be found in Bannai et al 2023^39^). In the second pass, brains were given to a new rater for further checking and editing, and a final evaluation was completed by an expert rater (DB) to ensure cohesion of manual segmentation across all raters. DB was trained by PL, a senior neuropsychiatrist with 8 years of clinical and image processing experience. The trigonum collaterale was used as a reliable starting point for segmentation. Manual segmentation started in the axial>sagittal>coronal views to ensure the entirety of the ChP was captured and that voxels from surrounding brain structures (e.g. fornix, hippocampus, corpus collosum, and thalamus) were excluded. A 3D inspection was performed at the final stage to ensure proper overall segmentation (Figure 1A-C). Third and fourth ventricle ChP were excluded to better compare our segmentation results with FreeSurfer and GMM, both of which only segment lateral ventricle ChP. ChP volume measures were extracted using ‘fslstats’ (FMRIB Software Library, v6.0). FreeSurfer was used to extract intracranial volume (ICV), lateral ventricle volume (LVV), total and subcortical gray matter volumes (GMV), and total white matter volume (WMV). Cortical measures greater than 4 standard deviations (SD) were removed (N=8) and those between 3-4 SD (N=21) were winsorized to the 3^rd^ SD. Manual segmentation was considered the gold standard with respect to FreeSurfer 6.0 and GMM segmentations of the lateral ventricle ChP.

**Figure 1.**
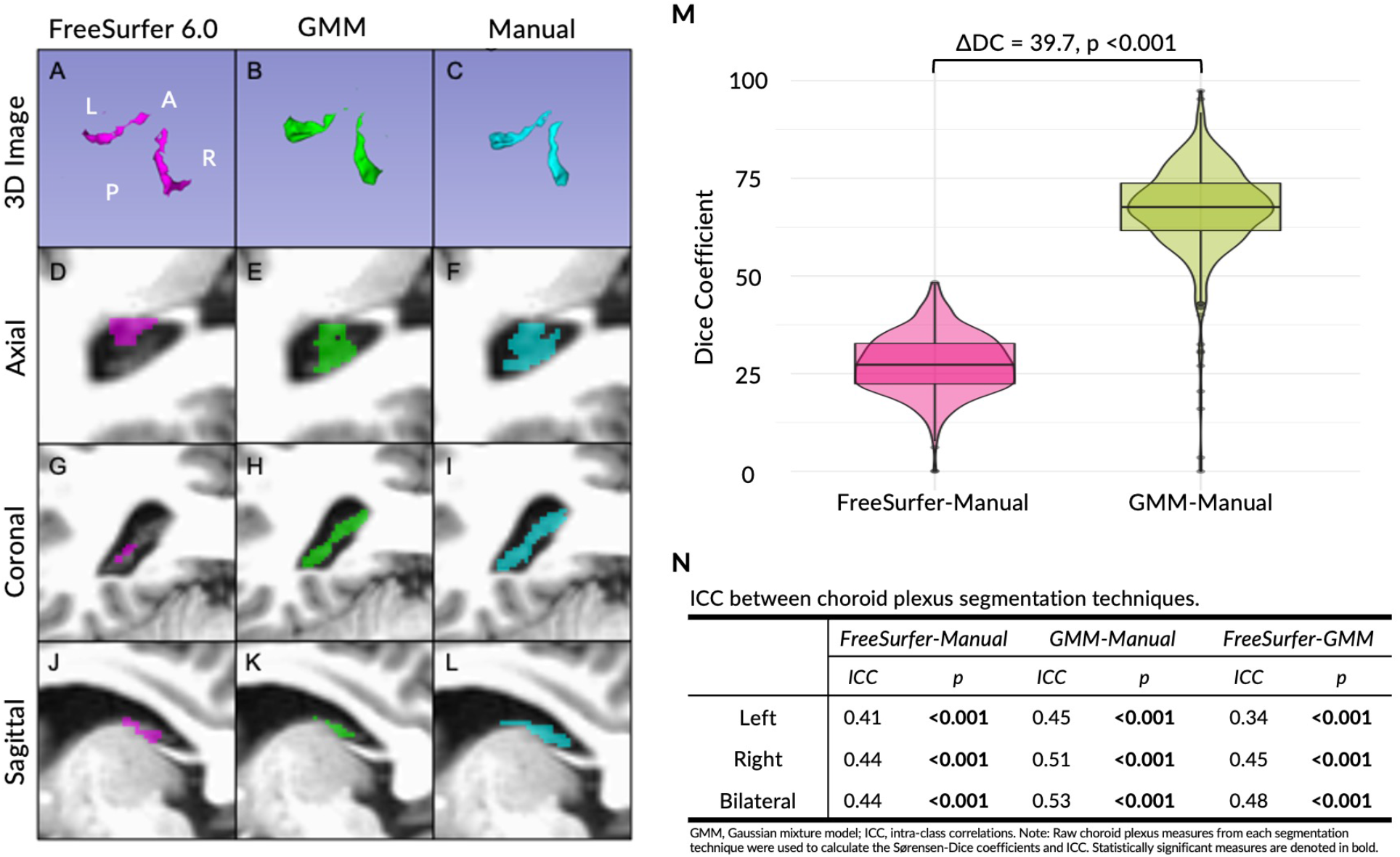
Schematic showing differences in choroid plexus segmentation between FreeSurfer 6.0 (Figures 1A, 1D, 1G, and 1J), Gaussian Mixture Model (Figures 1B, 1E, 1H, and 1K), and manual segmentation (Figures 1C, 1F, 1I, and 1L). White letters stand for: “A”, anterior; “P”, posterior”; “R”, right; “L”, left. Panel M demonstrates the difference in segmentation accuracy, quantified using Dice coefficients. Panel N enumerates intra-class correlation values across the three segmentation techniques.

### Plasma analyte collection

84 participants (33 HC and 51 CHR) had both MRI and plasma analyte data. A comprehensive detail of blood collection/assay has been provided previously^40^. In brief, blood was collected at baseline, processed within 120min and stored at -80°C. 185 analytes were immunoassayed blind to the subject’s clinical status. As previously described, 44 analytes were excluded due to medication or substance use confounds^40^. 18 analytes were not detected in >25% of subjects and were excluded. CHR vs HC comparisons were performed on the remaining 123 analytes and significant analytes were used for subsequent analyses.

### Statistical Analysis

Sociodemographic variables were compared between groups via Chi-square or analysis of variance (ANOVA) tests for categorical and continuous data, respectively. ANOVAs were performed to test the moderating effects of age, sex, race, site, ICV, and total GMV on manually-segmented ChP volume. Significant moderators (p<0.05) were used as covariates in ChP analysis and included age, sex, race, and site (Supplementary Table 1). ICV was used as an additional covariate for ChP measures. DC^41^, average Hausdorff distance^42^ (avgHD), and absolute intraclass correlation^43^ (ICC) were used to measure the accuracy and reliability of ChP segmentation between FreeSurfer, GMM, and manual segmentation. DC is a measure of the number of overlapping voxels segmented as ChP in two masks with values ranging from 0-100, where 0 designates no overlap and 100 reflects total overlap. Absolute ICC is a measure of agreement between two measurements and ranges from 0 to 1 (1=high reliability). General linear models, controlling for confounding variables, were used for group contrasts between CHR and HC. Post hoc comparisons examining conversion (converters vs non-converters), antipsychotic, duration (short vs long), and substance use status were performed. Effect sizes were calculated using Cohen’s d. To analyze whether ChP volume results were impacted by LVV or GMV, ChP group contrasts were performed using 1) LVV and 2) LVV and GMV in place of ICV as a covariate. Partial Spearman correlations were used to analyze associations between ChP volume and clinical (n= 10) and brain (n= 4). P-values were corrected for multiple comparisons using the false discovery rate (FDR) method (q-values)^44^. Finally, an exploratory analysis between ChP volume and blood analyte measures (n=123) was conducted using partial Spearman’s correlations in each diagnostic group (HC, n=33; CHR, n=51; Non-Converters, n=35; Converters, n=14). All statistical analyses were performed using R statistical software (version 3.5.1).

## Results

### Demographics

The final sample included 340 CHR and 169 HC (Table 1). HC and CHR demonstrated no race differences, but differed for age *(p=0*.*003)*, sex *(p=0*.*025)*, and site *(p=0*.*006)*. CHR had higher rates of substance use disorder (*p=0*.*004)*, SOPS (total, general, positive, negative, disorganization) and trauma scores *(p<0*.*001)*. CHR had lower GAF, MATRICS, and TASIT scores *(p<0*.*001)*. 105 (30%) CHR were on a first-or second-generation antipsychotic with an average daily chlorpromazine dosage of 187.8mg. In CHR with available data on conversion status (n=277), 36 (13%) converted to psychosis.

**Table 1.** Clinical and demographics information for HC and CHR groups.

| Variable | HC<br>(n=169) | CHR<br>(n=340) | Test Stat ($\chi^2, F$) | *p* |
| --- | --- | --- | --- | --- |
| Age (years) | 20.1 (4.8) | 18.9 (4.1) | 8.92 | **0.003** |
| Sex (Female/Male) | 87/82 | 138/202 | 5 | **0.025** |
| Race (CA/OT) | 93/76 | 192/147 | 0.062 | 0.81 |
| Site Number (1-8) | 20/16/26/20/23/16/23/25 | 38/45/23/37/55/33/79/30 | 19.6 | **0.006** |
| Antipsychotic Status (Yes/No) | - | 241/105 | - | - |
| CPZ Equivalent | - | 187.8 (216.6) | - | - |
| Converter/Non-Converter | - | 36/231 | - | - |
| Substance Use Disorder (Yes/No) | 1/168 | 24/332 | 8.25 | **0.004** |
| Duration of Illness (weeks) | - | 51.4 (75.7) | - | - |
| SOPS Total Score | 4.29 (5.99) | 37.6 (13.0) | 1010 | **<0.001** |
| SOPS General | 1.29 (2.25) | 8.84 (4.4) | 438 | **<0.001** |
| SOPS Positive | 0.94 (1.57) | 12.0 (4.2) | 1110 | **<0.001** |
| SOPS Negative | 1.43 (2.36) | 11.6 (6.2) | 416 | **<0.001** |
| SOPS Disorganization | 0.62 (1.15) | 5.19 (3.3) | 307 | **<0.001** |
| GAF | 84.12 (10.51) | 48.9 (11.4) | 1140 | **<0.001** |
| MATRICES Composite | 432.5 (65.1) | 391.2 (71.0) | 36.8 | **<0.001** |
| Social Cognition (TASIT) | 54.7 (5.4) | 52.9 (6.0) | 10.2 | **0.002** |
| Documentation of Trauma | 5.56 (2.5) | 7.41 (2.7) | 56.3 | **<0.001** |

**Note:** HC, healthy controls; CHR, clinical high risk for psychosis; CA, Caucasian; OT, other; Sites 1-8 are as follows: University of California Los Angeles, Emory, Harvard, Zucker Hillside Hospital, University of North Carolina, University of California San Diego, University of Calgary, and Yale. CPZ, chlorpromazine; SOPS, Scale of Prodromal Symptoms; GAF, Global Assessment of Functioning; MATRICES, Measurement and Treatment Research to Improve Cognition in Schizophrenia; TASIT, The Awareness of Social Inference Test. Statistical significance is denoted in bold.

### Accuracy and reliability of ChP segmentation

To ensure high intra-rater accuracy and reliability of the ChP, the final rater DB re-edited a randomized selection of 20 brains and demonstrated high accuracy (Bilateral DC=0.89 and avgHD=3.27mm^3^; Right DC=0.89 and avgHD=3.0mm^3^; Left DC=0.89 and avgHD=3.53mm^3^) and reliability (Bilateral ICC=0.97, p<0.001; Right ICC=0.96, p<0.001; Left ICC=0.98, p<0.001, Supplementary Figure 1). Compared to manual segmentation, FreeSurfer *(DC=29; left ICC=0*.*41, p<0*.*001; right ICC=0*.*44, p<0*.*001; bilateral ICC=0*.*44, p<0*.*001;* Figure 1M,N) and GMM *(DC=67; left ICC=0*.*45, p<0*.*001; right DC=0*.*51, p<0*.*001; bilateral ICC=0*.*53, p<0*.*001)* demonstrated poor accuracy and reliability. Compared to FreeSurfer, GMM demonstrated significantly higher accuracy (*ΔDC=0*.*37, p<0*.*001*). Thus, manually segmented ChP volumes were used for primary analyses.

### ChP differences

CHR demonstrated higher ChP volume compared to HC *(left: d=0*.*23, p=0*.*019, q=0*.*021; right: d=0*.*22, p=0*.*021, q=0*.*021; bilateral: d=0*.*23, p=0*.*017, q=0*.*021;* Figure 2A, Supplementary Table 2). This effect did not hold after controlling for LVV or LVV+GMV (Supplementary Table 2). For group differences using FreeSurfer or GMM ChP measures see Supplementary Table 2. In CHR, post hoc comparisons were performed using conversion (36 converters, 229 non-converters), antipsychotic (105 antipsychotics, 244 no antipsychotics), illness duration (155 short, 155 long), and substance use statuses (21 substance use, 323 no substance use; Supplementary Table 3). Compared to HC, non-converters had higher ChP volumes *(d=0*.*22, p=0*.*03*, Figure 2B; Supplementary Table 4). CHR on antipsychotics demonstrated larger ChP compared to HC *(d=0*.*26, p=0*.*035;* Figure 2C). Interestingly, CHR with shorter illness durations had larger ChP compared to HC *(d=0*.*27, p=0*.*018;* Figure 2D). Lastly, CHR both with and without a history of substance abuse had significantly higher ChP volumes compared to HC *(no-substance use d=0*.*20, p=0*.*04; substance use: d=0*.*54, p=0*.*017;* Figure 2E).

**Figure 2.**
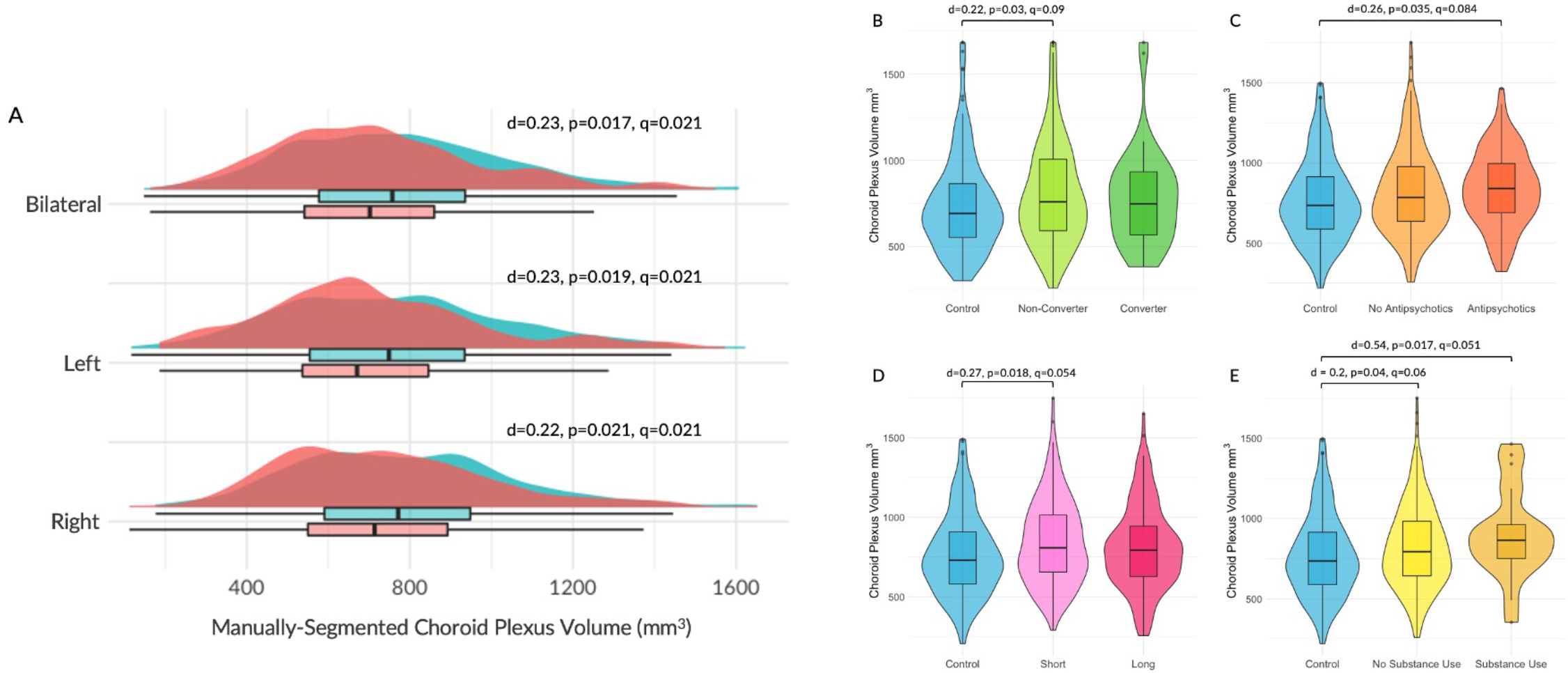
Panel A denotes diagnostic group differences between controls and CHR. Choroid plexus volume was corrected for age, sex, race, site number, and intracranial volume. Panels B-E demonstrate subgroup differences between controls and (B) non-converters and converters, (C) individuals on and not on antipsychotics, (D) individuals with short and long illness durations, and (E) individuals with and without a history of substance use. Note: ChP measures were covaried for age, sex, race, site number, and total intracranial volume.

### Clinical correlations

No significant correlations were observed between ChP volume and SOPS, SOPS subscales, global functioning, duration of illness, MATRICS, or trauma scores in the overall CHR, non-converter, or HC groups (Supplementary Table 5 and 6). In converters, greater ChP volume was associated with lower social cognition (*r=-0*.*37, p=0*.*031;* Supplementary Table 6). When examining the relationship between specific SOPS subscale scores and ChP volume, greater ChP volume was associated with poorer social anhedonia scores in CHR (*r=0*.*14, p=0*.*011*) and in non-converters (*r=0*.*14, p=0*.*032;* Supplementary Table 7). In converters, larger ChP volume correlated with lower experience of emotions and self (*r=0*.*39, p=0*.*019*), lower ideational richness (*r=0*.*34, p=0*.*045*), and lower amounts of grandiose ideas (*r=-0*.*41, p=0*.*013*; Supplementary Table 8). In CHR (overall, non-converters, and converters), no associations were noted between ChP volume and illness duration or CPZ equivalence (Supplementary Table 6).

### Neuroimaging correlations

In CHR, larger ChP was associated with lower total GMV *(r=-0*.*22, p<0*.*001, q<0*.*001)*, subcortical GMV *(d=-0*.*21, p<0*.*001, q<0*.*001)*, and WMV *(r=-0*.*28, p<0*.*001, q<0*.*001)*, as well as with higher LVV *(r=0*.*63, p<0*.*001, q<0*.*001;* Figure 3, Supplementary Table 9). HC demonstrated significant correlations between higher ChP and lower WMV *(r=-0*.*27, p<0*.*001, q<0*.*001)* and higher LVV *(r=0*.*66, p<0*.*001, q<0*.*001)*.

**Figure 3.**
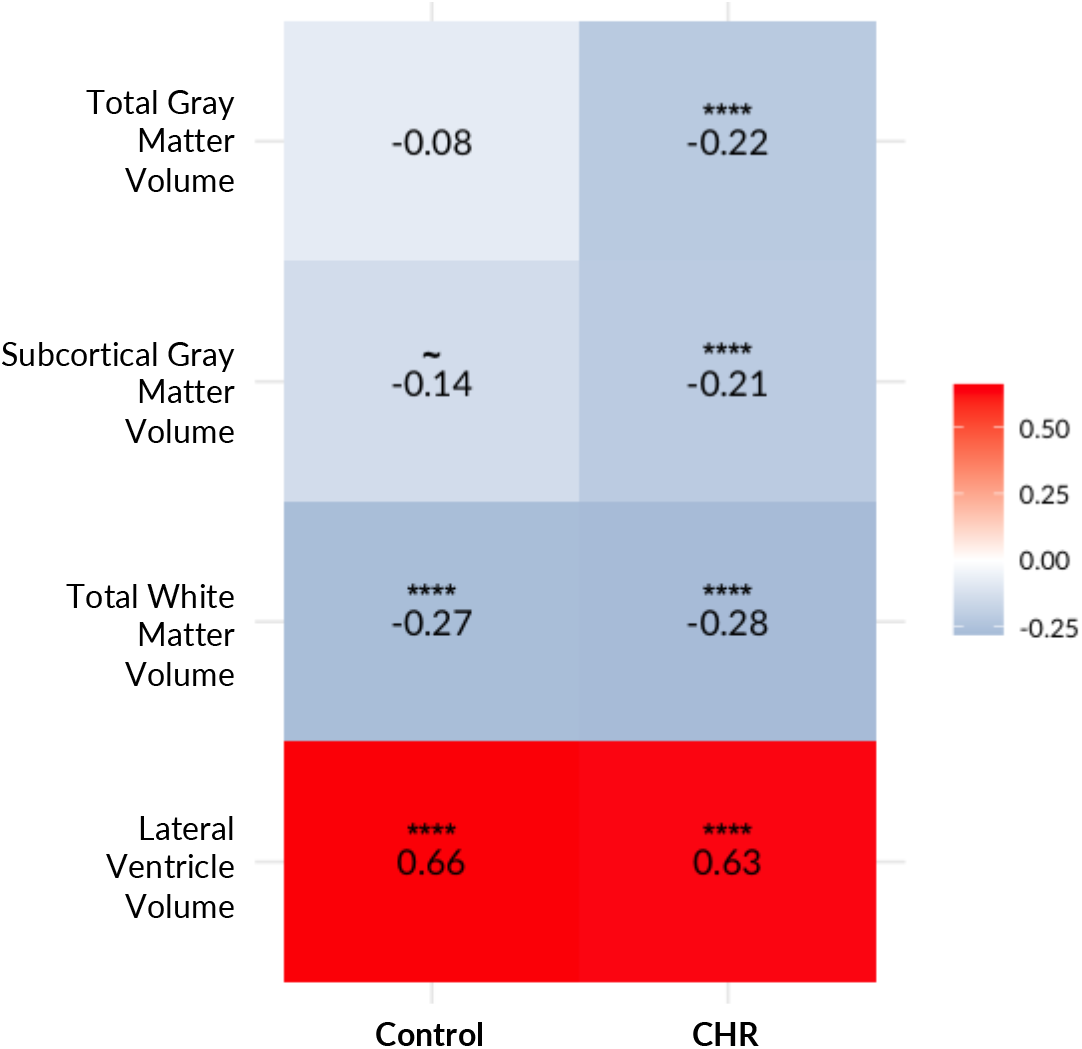
Choroid plexus volume correlations with cortical measures. Test statistics are denoted as ∼ p<0.1 and **** p<0.001.

**Figure 4.**
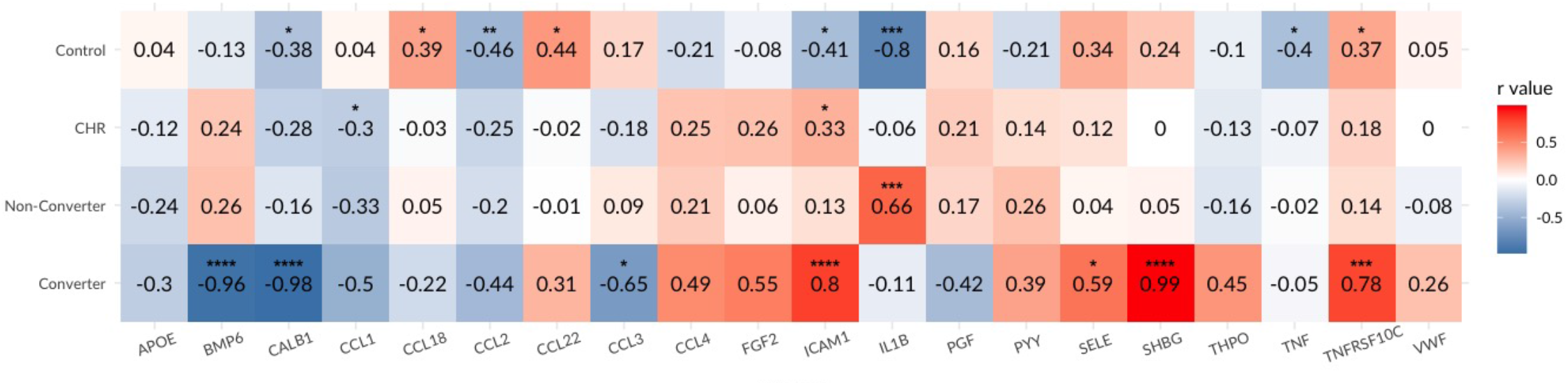
Exploratory correlational analysis between blood analyte measures and ChP volumes in each diagnostic group. Test statistics are denoted as * p<0.05, ** p<0.01, *** p<0.005, and **** p<0.001.

### Exploratory plasma analyte correlations

There were no sociodemographic differences between HC and CHR in the subset of participants with both MRI and plasma analyte data (Supplementary Table 10). A subset of analytes were chosen for exploratory correlational analyses based on the relative strength (*p* < 0.1) of their functional association with the lateral ventricle ChP (Harmonizome^45^) which resulted in 20 analytes being carried forward for further analyses.

In CHR, greater ChP volume was associated with higher levels of intracellular adhesion molecule 1 (ICAM1; *r=0*.*33, p=0*.*02)* and lower levels of lymphocyte secreted protein I-309 (CCL1; *r=-0*.*30, p=0*.*035*). Neither correlation remained significant after FDR-correction (Supplementary Table 11).

In controls, ChP volume was associated with higher chemokine ligand 18 (CCL18; *r=0*.*39, p=0*.*024*), chemokine ligand 22 (CCL22; *r=0*.*44, p=0*.*012*), and TNFRSF10C (*r=0*.*37, p=0*.*033*) levels as well as with lower serum CALB1 (*r=-0*.*38, p=0*.*031*), chemokine ligand 2 (CCL2; *r=-0*.*46, p=0*.*008)*, ICAM1 (*r=-0*.*41, p=0*.*019*), IL1β (*r=-0*.*80, p=0*.*003*), and tumor necrosis factor (TNF; *r=-0*.*40, p=0*.*032*; Supplementary Table 11). No associations survived FDR correction.

Non-converters demonstrated a significant positive relationship between ChP volume and peripheral interleukin 1β (IL1β; *r=0*.*66, p=0*.*004)*. This did not remain significant after FDR correction (Supplementary Table 12). Converters showed strong correlations between larger ChP volume and greater levels of ICAM1 (*r=0*.*80, p<0*.*001)*, E-selectin (SELE; *r=0*.*59, p=0*.*026*), sex hormone-binding globulin (SHBG; *r=0*.*99, p<0*.*001*), and tumor necrosis factor receptor superfamily member 10C (TNFRSF10C; *r=0*.*78, p<0*.*001*). Additionally, ChP volume in converters was negatively correlated with bone morphogenetic protein 6 (BMP6; *r=-0*.*96, p<0*.*001*), calbindin (CALB1; *r=-0*.*98, p<0*.*001*), and chemokine (C-C motif) ligand 3 (CCL3; *r=-0*.*65, p=0*.*017*; Supplementary Table 12). Of these, associations with BMP6 (*q<0*.*001*), CALB1 (*q<0*.*001*), ICAM1 (*q=0*.*003*), SHBG (*q<0*.*001*), and TNFRSF10C (*q=0*.*004*) remained significant after FDR correction.

## Discussion

Our study demonstrated that FreeSurfer and GMM had poor accuracy and reliability for the segmentation of the ChP compared to manual segmentation, which suggests that until a better automated tool exists that manual segmentation be undertaken when assessing ChP volume. We also showed that patients with CHR had larger ChP volume compared to HC. Moreover, patients that either did not convert to psychosis, had a shorter duration of illness, were on antipsychotics, or were using illicit substances had larger ChP volume compared to HC, with substance use demonstrating the largest effect. While no clinical or cognitive correlations were observed in the overall CHR group, converters demonstrated that larger ChP was associated with poorer social cognitive performance. It was also determined that greater ChP volume was associated with lower cortical and subcortical volumes, and larger LVV. Lastly, greater ChP volume correlated with makers functionally associated with the lateral ventricle ChP in CHR (CCL1, ICAM1), non-converters (IL1β), and converters (BMP6, CALB1, ICAM1, SELE, SHBG, TNFRSF10C). Together, these results suggest that the ChP may be an important explanatory biomarker for CHR.

The magnitude of ChP volume difference identified between CHR and HC in our study is of high significance since the effect size is similar to what has been reported by the ENIGMA consortium for cortical thickness measures (Cohen’s d ranging from -0.17 to -0.09)^46^ and because these changes are being identified prior to a conversion to psychosis or other psychiatric disorder has taken place. Additionally, these results extend upon prior findings of lateral ventricle ChP enlargement in first-episode^22^ and chronic psychosis^19^ to a CHR for psychosis population using a “gold standard” method for ChP segmentation. We expanded on the existing literature by demonstrating that ChP volume may be increased in individuals with shorter durations of illness, which hasn’t been demonstrated previously, but would have been predicted since prior studies have identified larger ChP volumes in early course^22^ and chronic psychosis^19^. We also found that antipsychotics were associated with greater ChP volume, which is inconsistent with our prior findings in early course^22^ and chronic psychosis^19^. Moreover, ChP enlargement has been reported in other neuropsychiatric disorders, such as Alzheimer’s^12^, multiple sclerosis (MS)^47,48^, neuromyelitis optica^47^, and major depressive disorder^13^. While these studies are promising, a reference for normal size criterion for the ChP is needed. Thus far, one study in children (0-16 years)^5^ and two studies in adults (18-94 years)^6,7^ have attempted to do this, but were limited by small sample sizes, measurement types (thickness vs volume), neuroimaging modality (MRI vs ultrasound), narrow age categories (mostly older adults), and use of less accurate ChP segmentation algorithms, which make it challenging to understand the role of ChP morphology in neuropsychiatric disorders. Nevertheless, the findings herein identified a novel and potentially important explanatory biomarker for CHR.

While no relationship was identified between ChP volume and overall cognitive performance in this study, these relationships have been described in case reports^49–51^ and in patients with psychosis spectrum disorders^19^. However, a significant relationship was identified in this study between greater ChP volume and lower performance on social cognitive testing in CHR converters, but not in non-converters or HC, which is a novel observation. While no study has previously described this relationship in other neuropsychiatric disorders it is possible that larger ChP volume in CHR converters may increase task related activity in frontal, temporal, and cingulate cortices, as well as abnormal responses to neutral stimuli during emotional processing tasks^52^. Though these findings are intriguing further work is needed to replicate these findings and to better understand the importance of this relationship.

The meaning of ChP enlargement in patients with psychosis has yet to be determined, but recent studies have shed light on this matter. For example, Lizano et al. (2019)^19^ observed relationships between ChP enlargement and larger LVV and lower total GMV in psychotic disorders. In MS, ChP enlargement was associated with higher LVV, greater WM lesions, lower brain volume, and normal-appearing WMV^48^. In depression, researchers found that ChP enlargement correlated with markers of central inflammation, namely greater mitochondrial translocator protein (TSPO) expression within the anterior cingulate, prefrontal, and insular cortices^13^. ChP enlargement was also associated with higher neuroinflammation in MS, reflected by greater ^18^F-DPA-714 binding in the thalamus^48^. The ChP of MS patients demonstrated greater TSPO expression compared to controls, suggesting an increased presence of immune cells within the ChP epithelium. Similarly, greater gadolinium enhancement of lateral and 4^th^ ventricle ChP in MS was identified suggesting a potential increase in ChP stromal capillary permeability^47^. Lastly, a recent study demonstrated that greater baseline ChP pT2 (pseudo T2, a measure ChP inflammation) was associated with clinical disability progression at follow-up^53^. These studies suggest that ChP enlargement is associated with inflammatory and microvascular changes within the ChP itself, as well as other parenchymal areas, which may hold the key for identifying explanatory biomarkers for disease progression.

An additional gap in the literature pertains to the effects of blood analytes on ChP morphology in psychosis. Previous work has demonstrated associations between ChP enlargement and higher peripheral IL6^19^ in those with psychotic disorders and with greater allostatic load (made up of cardiometabolic and C-reactive protein) in first-episode psychosis^22^. We were unable to examine IL6 in this study since it did not survive our quality control metrics. However, we found that enlarged ChP in CHR correlated with higher levels of ICAM1 and lower levels of CCL1, both of which are markers known to be functionally associated with the ChP^54–56^ and suggests the potential specificity of the relationships identified herein. The strongest and most consistent finding from our correlations related to ICAM1, which was found to be positively related to ChP volume in the overall CHR group and converters, not in the non-converters group, and orthogonal to the relationship observed in the HC group. ICAM1 is located on the apical surface of the ChP epithelium and its expression is increased with inflammatory activation. ICAM also supports leukocyte trafficking and immunosurveillance of the CNS, specifically at the ChP epithelium, which could potentially be a marker of CNS infalmmation^55^. The literature associating ICAM1 with psychosis has been mixed^57–59^. In addition to ICAM1, three other analytes (SELE, SHBG, and TNFRSF10C) positively correlated with ChP volume in converters. SELE functions similarly to ICAM1, can found on stromal venules in the ChP, and it also supports leukocyte trafficking and immunosurveillance of the CNS^55^. However, less is known about SELE in psychosis^57^. SHBG has been found in epithelial cells of the choroid plexus and ependymal cells of the third ventricle suggesting a role for neuroendocrine regulation^60^. Only one report identified increased levels of SHBG in antipsychotic naïve first episode psychosis male patients and it relevance to psychosis remains to be elucidated^61^. TNFRSF10C, a member of the TNF superfamily, is involved in apoptosis^62^ and no identifiable relationship to psychosis.

In the correlational analyses in CHR converters, greater ChP volume was associated with lower levels of CCL3, BMP6, and CALB1. CCL3 is robustly induced by the ChP during acute inflammation leading to the recruitment and activation of monocytes and neutrophils via the TLR2 receptor, a major receptor for gram positive bacteria, which also results in clefts and perforations of the basement membrane in ChP epithelial cells^63^. Despite this connection, few studies have described CCL3 differences in psychosis^64^. These results suggests that higher levels of CCL3 may result in lower ChP volume via ChP epithelial cell changes in CHR converters. BMP6, a ligand of the TGF superfamily, is involved in survival, neurite outgrowth, and is highly expressed in the ChP^65^. BMP6 expression is low in elderly schizophrenia patients^66^ and a BMP6 genetic locus on 6p24.3 was found to potentially play a role in impairments on sustained attention of schizophrenia^67^. CALB1, a member of the calcium-binding protein superfamily^68^, is expressed at low levels in the ChP^45^, and it expression was found to be decreased in post mortem brains of schizophrenia patients with high inflammation compared to low those with low inflammation^69^. While CALB1 expression was associated with ChP volume in CHR converters the relevance of this finding is unclear given the low expression of CALB1 in the ChP.

In non-converters, we found that enlarged ChP volume correlated with higher IL1β levels, which is orthogonal to the relationship observed in the HC group. IL1β is a proinflammatory cytokine that can be induced by stress^70^ and aging^71^. IL1β has been associated with the development of mood and psychotic symptoms^70^. In a recent mouse study, IL1β labeled macrophages were found in aged ChP which localized to endothelial, mesenchymal, and epithelial cells expressing IL1R1, which suggests that these interactions may contribute to age-dependent macrophage migration and infiltration of the ChP-brain immune barrier^71^. These finding align with the accelerating aging hypothesis in schizophrenia^72^ and other psychiatric disorders^73^, and suggests that CHR non-converters may be vulnerable to accelerated brain and cognitive changes.

While these analyte relationships are of great interest, what these studies don’t tell us is whether these markers are being released into the blood or CSF. It is possible that the ChP releases extracellular vesicles carrying inflammatory mediators to the blood or CNS^74^, but this hypothesis remains to be tested in psychosis. Nevertheless, the ChP is likely playing the role of immune surveillance, however, it is currently unknown whether the marker signature identified in CHR and converters is a result of an inherent deficit within ChP epithelium or a response to changes in the periphery or CNS.

In conclusion, our results suggest that the ChP is a biomarker of particular interest for neurodevelopment and CHR for psychosis. Strengths of this study include the use of manually-segmented ChP volume in the largest sample size of CHR patients. Limitations include 1) a sample size for examining ChP to peripheral analyte relationships, 2) lack of 3^rd^/4^th^ ventricle ChP segmentation given their proximity to important brain areas (hippocampus and cerebellum), and 3) cross-sectional neuroimaging design. Future studies should focus on these remaining gaps to would allow for a more compressive understanding of ChP morphology in psychosis.

## Acknowledgements

The authors would like to acknowledge Ehsan Tadayon for his help regarding choroid plexus segmentation techniques and for use of his Gaussian Mixture Model and the late Dr. Larry J. Seidman for his contribution to the field of CHR for psychosis research. The authors thank the participants at each recruitment site who took part in this study.

