## Supplementary Tables and Figures for "Linking Enlarged Choroid Plexus with Plasma Analyte and Structural Phenotypes in Clinical High Risk for Psychosis: A Multisite Neuroimaging Study"

**Supplementary Table 1.** Effects of moderators on mean ChP volume.

| Variable | *F* | *p* |
|---|---|---|
| Age | 1.01 | 0.32 |
| Sex | 47.1 | **<0.001** |
| Race | 0.038 | 0.85 |
| Site Number | 4.51 | **<0.001** |
| ICV | 51.7 | **<0.001** |
| Total GM Volume | 16.9 | **<0.001** |
| LV Volume | 265.7 | **<0.001** |
| Antipsychotic Status | 0.33 | 0.57 |
| CPZ Equivalent | 4.47 | **0.037** |

**Note:** ChP, choroid plexus; ICV, intracranial volume; LV, lateral ventricle; CPZ, chlorpromazine. Moderator analyses for antipsychotic status and chlorpromazine equivalents were conducted only within the prodromal sample. Statistical significance is denoted in bold.

**Supplementary Figure 1.** Schematic demonstrating choroid plexus segmentations by final rater DB for a first (Figures 1A, 1C, 1E, and 1G) and second (Figures 1B, 1D, 1F, and 1H) rating.  White letters stand for: “A”, anterior; “P”, posterior”; “R”, right; “L”, left. Panel I enumerates segmentation accuracy and reliability via intraclass correlation, Dice coefficient, and average Hausdorff distance.

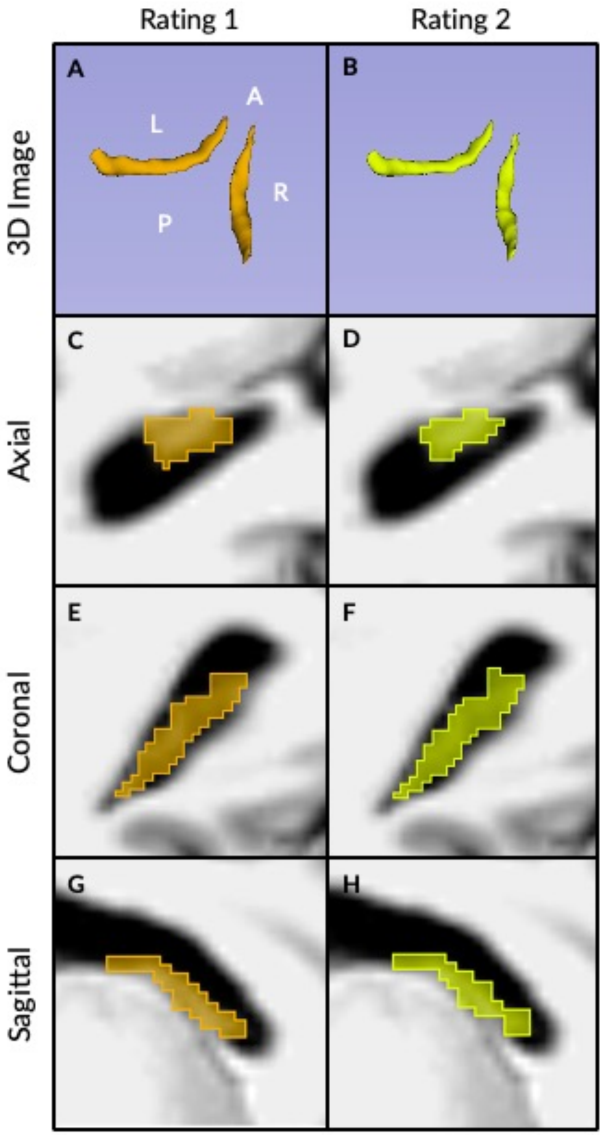


I

Quantitative measures of inter-rater accuracy and reliability for final rater DB.

| | | *Value* | *p* |
|---|---|---|---|
| ICC | Bilateral | 0.97 | **<0.001** |
| | Left | 0.98 | **<0.001** |
| | Right | 0.96 | **<0.001** |
| DC | Bilateral | 0.89 | - |
| | Left | 0.89 | - |
| | Right | 0.89 | - |
| avgHD | Bilateral | 3.27 $mm^3$ | - |
| | Left | 3.53 $mm^3$ | - |
| | Right | 3.0 $mm^3$ | - |

ICC, intraclass correlation; DC, Dice coefficient; avgHD, average Hausdorff distance. Note: Raw choroid plexus measures were used to calculate accuracy and reliability metrics. Statistically significant values are denoted in bold.

**Supplementary Table 2.** Group differences in ChP volume between three segmentation techniques.

| | *Covariates* | | *+ LVV* | | *+ LVV, GMV* | |
|---|---|---|---|---|---|---|
| Measure | *Cohen's d* | *p* | *Cohen's d* | *p* | *Cohen's d* | *p* |
| ***Freesurfer 6.0*** | | | | | | |
| Left Choroid Plexus | 0.26 | **0.004** | 0.14 | 0.12 | 0.12 | 0.18 |
| Right Choroid Plexus | 0.30 | **<0.001** | 0.25 | **0.006** | 0.22 | **0.016** |
| Overall Choroid Plexus | 0.32 | **<0.001** | 0.23 | **0.013** | 0.20 | **0.026** |
| ***Gaussian Mixture Model*** | | | | | | |
| Left Choroid Plexus | 0.24 | **0.008** | 0.13 | 0.14 | 0.14 | 0.12 |
| Right Choroid Plexus | 0.16 | 0.077 | 0.081 | 0.37 | 0.10 | 0.25 |
| Overall Choroid Plexus | 0.24 | **0.008** | 0.13 | 0.16 | 0.14 | 0.11 |
| ***Manual Segmentation*** | | | | | | |
| Left Choroid Plexus | 0.23 | **0.019** | 0.07 | 0.44 | 0.056 | 0.53 |
| Right Choroid Plexus | 0.22 | **0.021** | 0.15 | 0.082 | 0.11 | 0.23 |
| Overall Choroid Plexus | 0.23 | **0.017** | 0.082 | 0.37 | 0.066 | 0.46 |

**Note:** ChP, choroid plexus; LV, lateral ventricle; GM, gray matter. Choroid plexus measures were log transformed and covaried for age, sex, race, site number, and intracranial volume. Additional covariates, such as lateral ventricle volume and total gray matter volume, were added in the subsequent analyses demonstrated. Statistical significance is denoted in bold.

**Supplementary Table 3.** Clinical and demographics information for CHR converter, antipsychotic, illness duration, and substance use subgroups.

| Variable | **Non-Converters (n=229)** | **Converters (n=36)** | **Test Stat ($\chi^2$,*F*)** | ***p*** |
|---|---|---|---|---|
| Age (Years) | 18.9 (4.0) | 18.8 (4.0) | 0.04 | 0.84 |
| Sex (F/M) | 92/137 | 15/21 | <0.001 | 1 |
| Race (CA/OT) | 130/99 | 14/22 | 3.32 | 0.068 |
| Site Number | 23/31/18/30/38/22/48/19 | 4/7/3/1/9/2/7/3 | 5.47 | 0.60 |
| | **No Antipsychotics (n=244)** | **Antipsychotics (n=105)** | **Test Stat ($\chi^2$,*F*)** | ***p*** |
| Age (Years) | 19.1 (4.3) | 18.5 (3.7) | 1.57 | 0.21 |
| Sex (F/M) | 99/140 | 41/64 | 0.029 | 0.77 |
| Race (CA/OT) | 127/111 | 66/39 | 2.3 | 0.13 |
| Site Number | 21/38/14/21/35/24/63/23 | 17/8/9/16/20/10/16/9 | 15.9 | **0.026** |
| | **Short (n=155)** | **Long (n=155)** | **Test Stat ($\chi^2$,*F*)** | ***p*** |
| Age (Years) | 19.0 (4.1) | 18.9 (4.1) | 0.004 | 0.95 |
| Sex (F/M) | 61/94 | 67/88 | 0.33 | 0.56 |
| Race (CA/OT) | 85/70 | 89/65 | 0.17 | 0.68 |
| Site Number | 20/18/13/21/19/17/35/12 | 12/23/8/11/32/12/37/20 | 13.2 | 0.068 |
| | **No Substance Use (n=323)** | **Substance Use (n=21)** | **Test Stat ($\chi^2$,*F*)** | ***p*** |
| Age (Years) | 18.8 (4.1) | 20.7 (4.3) | 4.0 | **0.046** |
| Sex (F/M) | 132/191 | 8/13 | <0.001 | 0.98 |
| Race (CA/OT) | 180/142 | 13/8 | 0.10 | 0.76 |
| Site Number | 36/42/21/35/54/32/74/29 | 2/4/2/2/1/2/5/2 | 3.3 | 0.8 |

**Note:** CHR, clinical high-risk; HC, healthy controls; F, female; M, male; CA, Caucasian; OT, other; Statistical significance is denoted in bold.

**Supplementary Table 4.** Group differences in mean ChP volumes between HC and CHR, converter, medication, and duration subgroups.

| | *Cohen's d* | *p* |
|---|---|---|
| **Psychosis Conversion** | | |
| Control vs. Non-Converter | 0.22 | **0.03** |
| Control vs. Converter | 0.10 | 0.57 |
| Non-Converter vs. Converter | -0.10 | 0.57 |
| **Antipsychotic Usage** | | |
| Control vs. NonAP | 0.29 | 0.056 |
| Control vs. AP | 0.26 | **0.035** |
| NonAP vs. AP | 0.084 | 0.48 |
| **Duration** | | |
| Control vs. Short | 0.27 | **0.018** |
| Control vs. Long | 0.17 | 0.14 |
| Short vs. Long | -0.13 | 0.26 |
| **Substance Use** | | |
| Control vs. Non-Substance | 0.20 | **0.04** |
| Control vs. Substance | 0.54 | **0.017** |
| Non-Substance vs. Substance | 0.28 | 0.21 |

**Note:** ChP, choroid plexus; NonAP, CHR not on antipsychotic medications; AP, CHR on antipsychotic medications. Choroid plexus measures were adjusted for age, sex, race, site, and intracranial volume. Duration (short vs. long) was determined using a median split. Statistical significance is denoted in bold.

**Supplementary Table 5.** Spearman's correlations between mean ChP volume and clinical measures in CHR and HC.

| Clinical Measure | CHR | | HC | |
|---|---|---|---|---|
| | *r* | *p* | *r* | *p* |
| SOPS | 0.031 | 0.57 | -0.036 | 0.64 |
| SOPS General | 0.036 | 0.52 | -0.044 | 0.57 |
| SOPS Positive | -0.009 | 0.87 | -0.013 | 0.87 |
| SOPS Negative | 0.059 | 0.29 | 0.052 | 0.51 |
| SOPS Disorganization | 0.037 | 0.50 | 0.009 | 0.90 |
| Duration of Illness | -0.012 | 0.84 | - | - |
| GAF | -0.074 | 0.18 | -0.11 | 0.14 |
| MATRICS | -0.095 | 0.095 | -0.041 | 0.61 |
| Trauma | 0.044 | 0.43 | 0.042 | 0.59 |
| Social Cognition | -0.042 | 0.45 | -0.057 | 0.47 |
| CPZ Equivalent | -0.11 | 0.30 | - | - |

**Note:** ChP, choroid plexus; CHR, clinical high-risk; HC, healthy controls; SOPS, Scale of Prodromal Symptoms; GAF, Global Assessment of Functioning; MATRICS, Measurement and Treatment Research to Improve Cognition in Schizophrenia; CPZ, chlorpromazine. Choroid plexus and clinical measures were covaried for age, sex, race, and site number. Choroid plexus measures were additionally covaried for intracranial volume.

**Supplementary Table 6.** Spearman's correlations between mean ChP volume and clinical measures in converters and non-converters.

| Clinical Measure | Converters | | Non-Converters | |
|---|---|---|---|---|
| | *r* | *p* | *r* | *p* |
| SOPS | 0.021 | 0.90 | 0.027 | 0.69 |
| SOPS General | 0.091 | 0.60 | 0.004 | 0.95 |
| SOPS Positive | -0.066 | 0.70 | 0.003 | 0.97 |
| SOPS Negative | 0.17 | 0.33 | 0.021 | 0.76 |
| SOPS Disorganization | -0.019 | 0.91 | 0.08 | 0.23 |
| Duration of Illness | 0.026 | 0.88 | 0.052 | 0.46 |
| GAF | -0.031 | 0.065 | -0.096 | 0.15 |
| MATRICS | -0.25 | 0.17 | -0.11 | 0.13 |
| Trauma | -0.069 | 0.69 | -0.018 | 0.79 |
| Social Cognition | -0.37 | **0.031** | 0.025 | 0.71 |
| CPZ Equivalent | 0.18 | 0.69 | -0.25 | 0.058 |

**Note:** ChP, choroid plexus; SOPS, Scale of Prodromal Symptoms; GAF, Global Assessment of Functioning; MATRICS, Measurement and Treatment Research to Improve Cognition in Schizophrenia; CPZ, chlorpromazine. Choroid plexus and clinical measures were covaried for age, sex, race, and site number. Choroid plexus measures were additionally covaried for intracranial volume. Statistical significance denoted in bold.

**Supplementary Table 7.** Spearman's correlations between mean ChP volume and clinical measures in CHR individuals and HC.

| SOPS | Clinical Measure | CHR | | HC | |
|---|---|---|---|---|---|
| | | *r* | *p* | *r* | *p* |
| G1 | Sleep Disturbance | 0.013 | 0.81 | -0.017 | 0.83 |
| G2 | Dysphoric Mood | -0.003 | 0.95 | -0.019 | 0.81 |
| G3 | Motor Disturbances | 0.031 | 0.57 | -0.037 | 0.63 |
| G4 | Impaired Tolerance to Normal Stress | 0.073 | 0.19 | 0.013 | 0.87 |
| P1 | Unusual Thought Content/Delusions | 0.004 | 0.94 | 0.045 | 0.56 |
| P2 | Suspiciousness/Persecutory Ideas | 0.023 | 0.68 | 0.03 | 0.70 |
| P3 | Grandiose Ideas | -0.04 | 0.47 | 0.12 | 0.13 |
| P4 | Perceptual Abnormalities/Hallucinations | -0.04 | 0.47 | -0.074 | 0.34 |
| P5 | Disorganized Communication | -0.046 | 0.40 | -0.08 | 0.30 |
| N1 | Social Anhedonia | 0.14 | **0.011** | -0.007 | 0.93 |
| N2 | Avolition | 0.019 | 0.74 | 0.096 | 0.22 |
| N3 | Decreased Emotional Expression | 0.39 | 0.48 | 0.049 | 0.53 |
| N4 | Decreased Experience of Emotions and Self | 0.023 | 0.68 | 0.018 | 0.82 |
| N5 | Decreased Ideational Richness | 0.025 | 0.64 | -0.061 | 0.43 |
| N6 | Occupational Functioning | 0.011 | 0.85 | 0.048 | 0.53 |
| D1 | Odd Behavior/Appearance | 0.057 | 0.30 | -0.029 | 0.71 |
| D2 | Bizarre Appearance | -0.02 | 0.67 | 0.093 | 0.23 |
| D3 | Trouble with Attention/Focus | -0.006 | 0.92 | 0.03 | 0.70 |
| D4 | Impairment in Personal Hygiene | -0.001 | 0.98 | 0.007 | 0.93 |

**Note:** ChP, choroid plexus; CHR, clinical high-risk; HC, healthy controls; SOPS, Scale of Prodromal Symptoms; GAF, Global Assessment of Functioning; MATRICS, Measurement and Treatment Research to Improve Cognition in Schizophrenia; CPZ, chlorpromazine. Choroid plexus and clinical measures were covaried for age, sex, race, and site number. Choroid plexus measures were additionally covaried for intracranial volume. Statistical significance is denoted in bold.

**Supplementary Table 8.** Spearman's correlations between mean ChP volume and clinical measures in converters and non-converters.

| SOPS | Clinical Measure | Converters | | Non-Converters | |
|---|---|---|---|---|---|
| | | *r* | *p* | *r* | *p* |
| G1 | Sleep Disturbance | 0.031 | 0.86 | -0.021 | 0.75 |
| G2 | Dysphoric Mood | 0.041 | 0.81 | -0.010 | 0.88 |
| G3 | Motor Disturbances | 0.32 | 0.06 | 0.012 | 0.86 |
| G4 | Impaired Tolerance to Normal Stress | 0.008 | 0.96 | 0.051 | 0.44 |
| P1 | Unusual Thought Content/Delusions | 0.000 | 1.00 | 0.036 | 0.59 |
| P2 | Suspiciousness/Persecutory Ideas | 0.017 | 0.92 | 0.010 | 0.88 |
| P3 | Grandiose Ideas | -0.41 | **0.013** | 0.014 | 0.83 |
| P4 | Perceptual Abnormalities/Hallucinations | 0.057 | 0.74 | -0.025 | 0.71 |
| P5 | Disorganized Communication | 0.22 | 0.20 | -0.083 | 0.21 |
| N1 | Social Anhedonia | 0.013 | 0.94 | 0.14 | **0.032** |
| N2 | Avolition | -0.18 | 0.29 | -0.008 | 0.91 |
| N3 | Decreased Emotional Expression | 0.20 | 0.25 | -0.003 | 0.96 |
| N4 | Decreased Experience of Emotions and Self | 0.39 | **0.019** | -0.028 | 0.67 |
| N5 | Decreased Ideational Richness | 0.34 | **0.045** | 0.035 | 0.60 |
| N6 | Occupational Functioning | 0.13 | 0.45 | -0.010 | 0.89 |
| D1 | Odd Behavior/Appearance | -0.050 | 0.77 | 0.089 | 0.18 |
| D2 | Bizarre Appearance | -0.10 | 0.56 | -0.001 | 0.99 |
| D3 | Trouble with Attention/Focus | -0.11 | 0.53 | 0.014 | 0.83 |
| D4 | Impairment in Personal Hygiene | -0.059 | 0.73 | 0.052 | 0.44 |

**Note:** ChP, choroid plexus; SOPS, Scale of Prodromal Symptoms; GAF, Global Assessment of Functioning; MATRICS, Measurement and Treatment Research to Improve Cognition in Schizophrenia; CPZ, chlorpromazine. Choroid plexus and clinical measures were covaried for age, sex, race, and site number. Choroid plexus measures were additionally covaried for intracranial volume. Statistical significance is denoted in bold.

**Supplementary Table 9.** Spearman's correlations between ChP volume and cortical measures in CHR and controls.

| Measure | CHR | | | HC | | |
|---|---|---|---|---|---|---|
| | *r* | *p* | *q* | *r* | *p* | *q* |
| Total GMV | -0.22 | **<0.001** | **<0.001** | -0.08 | 0.31 | 0.31 |
| Subcortical GMV | -0.21 | **<0.001** | **<0.001** | -0.14 | 0.062 | 0.083 |
| Total WMV | -0.28 | **<0.001** | **<0.001** | -0.27 | **<0.001** | **<0.001** |
| LVV | 0.63 | **<0.001** | **<0.001** | 0.66 | **<0.001** | **<0.001** |

**Note**: ChP, choroid plexus; CHR, clinical high-risk; HC, healthy control; GMV, gray matter volume; WMV, white matter volume; LVV, lateral ventricle volume. All measures were adjusted for age, sex, race, site, and total intracranial volume. Statistical significance is denoted in bold.

**Supplementary Table 10.** Clinical and demographic information within individuals with neuroimaging and cytokine data.

| Variable | HC (n=33) | CHR (n=51) | Test Stat ($\chi^2$,$F$) | *p* |
|---|---|---|---|---|
| Age (years) | 20.1 (4.7) | 19.4 (4.4) | 0.41 | 0.52 |
| Sex (Female/Male) | 12/21 | 20/31 | 0.001 | 0.97 |
| Race (CA/OT) | 19/14 | 26/25 | 0.14 | 0.71 |
| Site Number | 2/7/6/3/4/0/7/4 | 2/12/5/4/13/1/11/3 | 4.7 | 0.70 |
| Antipsychotic Status (Yes/No) | - | | - | - |
| CPZ Equivalent | - | | - | - |
| Converter/Non-Converter | - | 14/35 | - | - |
| Substance Use Disorder (Yes/No) | 0/33 | 2/49 | 0.18 | 0.68 |
| SOPS Score | 5.03 (5.2) | 39.94 (14.2) | 184 | **p<0.001** |
| SOPS Positive | 1.45 (1.9) | 13.25 (5.0) | 169 | **p<0.001** |
| SOPS Negative | 1.18 (1.8) | 12.37 (6.3) | 99.9 | **p<0.001** |
| SOPS Disorganization | 0.91 (1.2) | 6.18 (3.5) | 70.1 | **p<0.001** |
| GAF | 85.03 (7.6) | 47.2 (11.0) | 300 | **p<0.001** |
| MATRICS Composite | 428.14 (63.1) | 391.68 (73.1) | 5.03 | **0.028** |
| Documentation of Trauma | 5.91 (2.2) | 8.04 (2.4) | 17 | **p<0.001** |
| Social Cognition (TASIT) | 54.72 (5.2) | 54.15 (6.4) | 0.17 | 0.68 |

**Note**: HC, healthy controls; CHR, clinical high-risk; CA, Caucasian; OT, other; CPZ, chlorpromazine; SOPS, Scale of Prodromal Symptoms; GAF, Global Assessment of Functioning; MATRICS, Measurement and Treatment Research to Improve Cognition in Schizophrenia; BACS, Brief Assessment of Cognition in Schizophrenia; TASIT, The Awareness of Social Inference Test. Statistical significance is denoted in bold.

**Supplementary Table 11.** Spearman's correlations between mean ChP volume and cytokine levels within diagnostic groups.

| Cytokine | CHR (n=51) | | | HC (n=33) | | |
|---|---|---|---|---|---|---|
| | *r* | *p* | *q* | *r* | *p* | *q* |
| APOE | -0.12 | 0.39 | 0.57 | 0.042 | 0.82 | 0.84 |
| BMP6 | 0.24 | 0.096 | 0.32 | -0.13 | 0.47 | 0.62 |
| CALB1 | -0.28 | 0.055 | 0.32 | -0.38 | **0.031** | 0.084 |
| CCL1 | -0.30 | **0.035** | 0.32 | 0.038 | 0.84 | 0.84 |
| CCL18 | -0.035 | 0.81 | 0.95 | 0.39 | **0.024** | 0.084 |
| CCL2 | -0.25 | 0.081 | 0.32 | -0.46 | **0.008** | 0.078 |
| CCL22 | -0.015 | 0.92 | 0.99 | 0.44 | **0.012** | 0.078 |
| CCL3 | -0.18 | 0.23 | 0.50 | 0.17 | 0.36 | 0.55 |
| CCL4 | 0.25 | 0.074 | 0.32 | -0.21 | 0.24 | 0.42 |
| FGF2 | 0.26 | 0.18 | 0.50 | -0.084 | 0.76 | 0.84 |
| ICAM1 | 0.33 | **0.02** | 0.32 | -0.41 | **0.019** | 0.084 |
| IL1β | -0.057 | 0.78 | 0.95 | -0.80 | **0.003** | 0.058 |
| PGF | 0.21 | 0.29 | 0.57 | 0.16 | 0.46 | 0.62 |
| PYY | 0.14 | 0.34 | 0.57 | -0.21 | 0.25 | 0.42 |
| SELE | 0.12 | 0.40 | 0.57 | 0.34 | 0.055 | 0.12 |
| SHBG | 0.003 | 0.98 | 0.99 | 0.24 | 0.17 | 0.34 |
| THPO | -0.13 | 0.36 | 0.57 | -0.10 | 0.57 | 0.71 |
| TNF | -0.071 | 0.65 | 0.86 | -0.40 | **0.032** | 0.084 |
| TNFRSF10C | 0.18 | 0.21 | 0.50 | 0.37 | **0.033** | 0.084 |
| VWF | -0.002 | 0.99 | 0.99 | 0.052 | 0.77 | 0.84 |

**Note:** Choroid plexus (ChP) and clinical measures were covaried for age, sex, race, and site number. Choroid plexus measures were additionally covaried for intracranial volume. Statistical significance denoted in bold. p-values were adjusted using false discovery rate correction (q-value).

**Supplementary Table 12.** Spearman's correlations between mean ChP volume and cytokine levels within converters and non-converters.

| Cytokine | Non-Converters (n=35) | | | Converters (n=14) | | |
|---|---|---|---|---|---|---|
| | *r* | *p* | *q* | *r* | *p* | *q* |
| APOE | -0.24 | 0.16 | 0.63 | -0.30 | 0.30 | 0.40 |
| BMP6 | 0.26 | 0.13 | 0.63 | -0.96 | **<0.001** | **<0.001** |
| CALB1 | -0.16 | 0.37 | 0.81 | -0.98 | **<0.001** | **<0.001** |
| CCL1 | -0.33 | 0.056 | 0.56 | -0.50 | 0.071 | 0.16 |
| CCL18 | 0.052 | 0.76 | 0.92 | -0.22 | 0.44 | 0.49 |
| CCL2 | -0.20 | 0.24 | 0.68 | -0.44 | 0.11 | 0.21 |
| CCL22 | -0.014 | 0.94 | 0.94 | 0.31 | 0.28 | 0.40 |
| CCL3 | 0.086 | 0.63 | 0.92 | -0.65 | **0.017** | 0.056 |
| CCL4 | 0.21 | 0.21 | 0.68 | 0.49 | 0.074 | 0.16 |
| FGF2 | 0.064 | 0.80 | 0.92 | 0.55 | 0.20 | 0.31 |
| ICAM1 | 0.13 | 0.47 | 0.81 | 0.80 | **<0.001** | **0.003** |
| IL1β | 0.66 | **0.004** | 0.08 | -0.11 | 0.78 | 0.82 |
| PGF | 0.17 | 0.48 | 0.81 | -0.42 | 0.40 | 0.49 |
| PYY | 0.26 | 0.14 | 0.63 | 0.39 | 0.17 | 0.29 |
| SELE | 0.038 | 0.83 | 0.92 | 0.59 | **0.026** | 0.074 |
| SHBG | 0.050 | 0.78 | 0.92 | 0.99 | **<0.001** | **<0.001** |
| THPO | -0.16 | 0.37 | 0.81 | 0.45 | 0.11 | 0.21 |
| TNF | -0.015 | 0.93 | 0.94 | -0.049 | 0.89 | 0.89 |
| TNFRSF10C | 0.14 | 0.42 | 0.81 | 0.78 | **0.001** | **0.004** |
| VWF | -0.075 | 0.67 | 0.92 | 0.26 | 0.44 | 0.49 |

**Note:** Choroid plexus (ChP) and clinical measures were covaried for age, sex, race, and site number. Choroid plexus measures were additionally covaried for intracranial volume. Statistical significance denoted in bold. p values were adjusted using false discovery rate correction.
